# Differential human mobility and local variation in human infection attack rate

**DOI:** 10.1101/291674

**Authors:** D.J. Haw, D. A. T. Cummings, J. Lessler, H. Salje S, J. M. Read, S. Riley

## Abstract

Infectious disease transmission in animals is an inherently spatial process in which a host’s home location and their social mixing patterns are important, with the mixing of infectious individuals often different to that of susceptible individuals. Although incidence data for humans have traditionally been aggregated into low-resolution data sets, modern representative surveillance systems such as electronic hospital records generate high volume case data with precise home locations. Here, we use a high resolution gridded spatial transmission model of arbitrary resolution to investigate the theoretical relationship between population density, differential population movement and local variability in incidence. We show analytically that uniform local attack rate is only possible for individual pixels in the grid if susceptible and infectious individuals move in the same way. Using a population in Guangdong, China, for which a robust quantitative description of movement is available (a movement kernel), and a natural history consistent with pandemic influenza; we show that for the estimated kernel, local cumulative incidence is positively correlated with population density when susceptible individuals are more connected in space than infectious individuals. Conversely, when infectious individuals are more connected, local cumulative incidence is negatively correlated with population density. The amplitude of correlation is substantial for the estimated kernel. However, the strength and direction of correlation changes sign for other kernel parameter values. These results describe a precise relationship between the spatio-social mixing of infectious and susceptible individuals and local variability in attack rates, and suggest a plausible mechanism for the counter-intuitive scenario in which local incidence is lower on average in less dense populations. Also, these results suggest that if spatial transmission models are implemented at high resolution to investigate local disease dynamics, including micro-tuning of interventions, the underlying detailed assumptions about the mechanisms of transmission become more important than when similar studies are conducted at larger spatial scales.

**Author Summary:** We know that some places have higher rates of infectious disease than others. However, at the moment, we usually only measure these differences for large towns and cities. With modern data, such as those we can get from mobile phones, we can measure rates of infection at much smaller scales. In this paper, we used a computer simulation of an epidemic to propose ways that rates of incidence in small local areas might be related to population density. We found that if infectious people are better connected than non-infectious people, perhaps because they receive visitors, then, on average, higher density areas would have lower rates of infection. If infectious people were less connected than non-infectious people then higher density areas would have higher rates of infection. As data get more accurate, this type of analysis will allow us to propose and test ways to optimize interventions such as the delivery of vaccines and antivirals during a pandemic.

## Introduction

The spatial heterogeneity of infectious disease incidence at large scales presents numerous intervention opportunities and challenges. Maps of malaria prevalence [1] have been used to target additional surveillance and to prioritize countries and geographical regions for additional intervention investment, resulting in substantial decreases in numbers of infections [2]. Over shorter timescales, spatial asynchrony in the northern hemisphere during the 2009 influenza pandemic likely led to variable effectiveness of vaccination when eventually deployed because of prior infections [3]. The epidemiological implications of substantial spatial heterogeneity in both incidence and transmission are topics of active research for most human pathogens [4].

These spatial heterogeneities must be influenced by two key human behaviours: where people choose to live and how they move. Because the home location of an individual is primarily used as the geographic location when cases are recorded, absolute spatial incidence is driven by population density: where more people live in a given unit area, there is greater potential for cases. Accurate high resolution estimates of population density [5, 6] have helped refine global absolute estimates of disease incidence and prevalence [7, 8, 9]. In order for a directly transmitted human pathogen to move through space, at least one person must travel away from home and meet another person. Even for vector borne pathogens such as malaria and Zika virus, typical distances traveled by the vector are much shorter than those traveled by human hosts. Human movement is captured by survey data on journeys to work [10], questionnaire-based surveys [11] and location logging of mobile devices [12, 13, 14].

Although spatial heterogeneity has been measured mainly at larger scales, modern pathogen surveillance enables more finely resolved incidence data sets, with details such as precise geographical location captured with increasing frequency by modern digital and biological technology. For example, the full genome of a pathogen can be made available in almost real time directly from clinical samples taken in the community [15], and the home location of everyone attending a health care facility can be extracted from clinical episode data [16]. Because this level of geographical precision for high quality incidence data has not previously been available, both epidemiological and disease-dynamic studies of infectious disease have focused on predicting and explaining incidence patterns measured at larger spatial scales, often with all cases within an administrative unit reported together. Additional insights are likely being lost during this aggregation process.

Available evidence and intuition suggests that infectious and non-infectious individuals have different social interactions during an outbreak [17], with plausible scenarios in which either one or the other may be more connected in space. For example, susceptible individuals are more likely to travel than are infectious individuals with mild symptoms [18]. However, family members and friends of infectious individuals may often not behave in the same way as an average susceptible individual. Also, infectious individuals themselves may travel long distances away from transmission hotspots to seek medical care during outbreaks of highly pathogenic infections [19].

Disease dynamic models are often used to study infection incidence and are defined primarily by their force-of-infection (FOI) term: a precise mathematical specification of how the risk of infection experienced by a susceptible individual is driven by the number of currently infectious individuals and by their characteristics. For example, the ages of infectious and susceptible individuals must sometimes affect the risk of infection, as must the distance between their home addresses. Disease dynamic models that represent space [20] are now used routinely to understand large-scale spatial heterogeneity in incidence: to estimate the relative effectiveness of spatially heterogeneous interventions (given the observed incidence); to reveal underlying social mechanisms of transmissions; and, with increasing frequency, to forecast future spatial incidence patterns [21]. All transmission models that represent space include some kind of spatial kernel-a formal definition of the way in which individuals from different locations distribute their influence over the whole of geographical space.

However, there is substantial variability in the underlying FOI assumptions made in these models, which are often not discussed explicitly and have likely only rarely made material differences to model-based results aggregated at larger spatial scales. Nonetheless, we hypothesise that these different FOI assumptions represent important alternate hypotheses for the mechanisms of transmission and may lead to substantial structural biases in the predictions of attack rates at smaller spatial scales. Here, we propose a general theoretical framework for the study of infectious disease incidence at arbitrarily small spatial scales and, in particular, we look at the relative mobility of infectious individuals relative to susceptible individuals as a potential driver of heterogeneity in incidence.

## Results

Algebraic analyses show that differential spatial connectivity of susceptible and infectious individuals can lead to variability in local attack rates (Protocol S1). Firstly, we showed that if susceptible and infectious individuals are assumed to be connected in the same way across all points in space, then local attack rates are uniform for any population density distribution or grid resolution. For lower resolution grids with large individual spatial elements, where the amplitude of connectivity of individuals outside their home grid square is small, the impact of differential connectivity between susceptible and infectious individuals is still negligible, even to the point that it is reasonable to assume that infectious individuals have no connectivity at all outside their home location. However, as the resolution of the grid increases and squares become smaller, individuals have a substantial number of connections outside their own grid square. Under this scenario, it was no longer possible to prove analytically that differences in the connectedness of susceptible and infectious individuals would not lead to local variation in attack rates. These analytical results were not affected by the presence of age stratification in the transmission process, so long as the behavior and distribution of age groups was assumed to be uniform across space.

We established a baseline numerical scenario by implementing the underlying transmission model (see Methods) as ordinary differential equations (ODEs). Using: a 1 k^2^ gridded population density (55 by 33 k to the east and north of Guangzhou, China); a spatial contact kernel estimated in the same population [22]; and a basic reproductive number *R*_0_ = 1.8; we recovered a global uniform attack rate of *z* = 0.73. We also introduced age-stratified populations and transmission using parameters estimated in this population [11]. For this population, accurate high-resolution data on local age distributions were not available, therefore, we assumed that all grid squares had populations with the same age distribution, even though the total number of individuals in a grid square varied substantially. This addition of age effects in the transmission process did not introduce spatial variation but did reduced the uniform global attack rate to *z* = 0.43. We validated the precision of attack rates obtained from the ODEs using age- and space-stratified refinements [20] of the standard implicit equation relating attack rate to *R*_0_ *z* = 1 *− e*^*−R*_0_*z*^[23] and.

We hypothesized that both population density and the gradient of population density may influence small-scale attack rates in these models. Figures 1A and 1B show the uniform attack rate when using dual mobility with four age classes, plotted against log of population density and gradient of log population density respectively (with log gradient defined as the average difference between the log of a location’s resident population and that of its 8 immediate neighbours).

**Figure 1:**
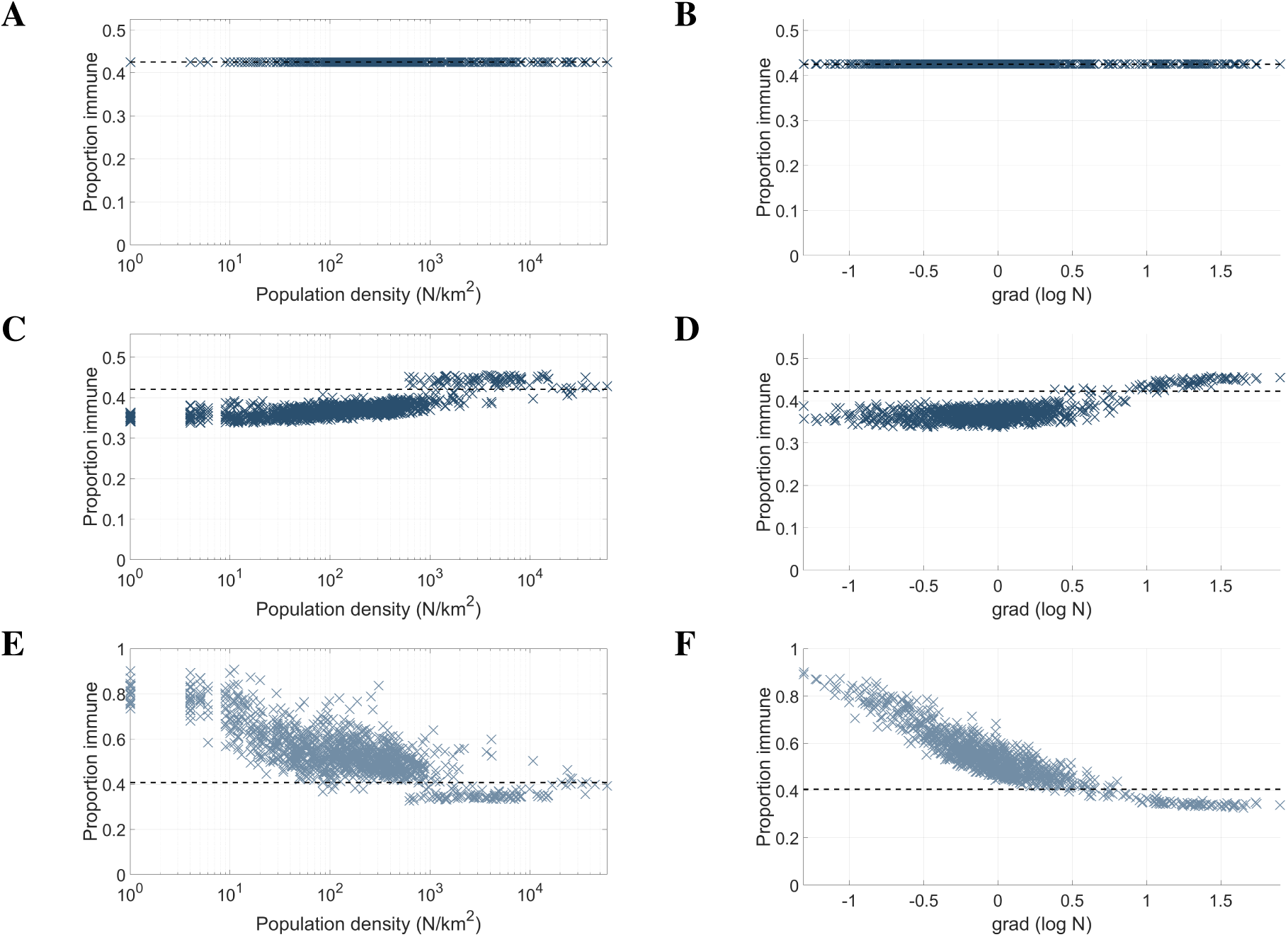
The relationship between: force-of-infection (FOI) assumptions, local attack rates, population density and population density gradient; for a pandemic-influenza-like epidemic. FOI assumptions are explained in the text. The LHS shows the relationship between the log10 population density and attack rate for **(A)** dual mobility, **(C)** S-mobility and **(E)** I-mobility. The RHS shows the relationship between the gradient of log10 population density and attack rate for **(B)** dual mobility, **(D)** S-mobility and **(F)** I-mobility. Population gradient was defined as the difference between the population density of a cell and the average population density of the 9 surrounding cells in the square lattice.

When only susceptible individuals were assumed to be mobile, location-specific attack rates were positively correlated with log population density, correlation coefficient c=0.75 (Figure 1C). Attack rates varied between a minimum of 33.72% to a maximum of 45.76%, an absolute range of 12.04%. Location-specific attack rates were slightly less correlated with the log gradient of population density (correlation coefficient c=0.73, Figure 1D). Locations with higher attack rates tended to be densely populated relative to neighboring locations (Figures 2A and 2C). Figure 2B illustrates the uniformity in attack rate obtained from dual mobility (locations with attack rate 0 are empty).

**Figure 2:**
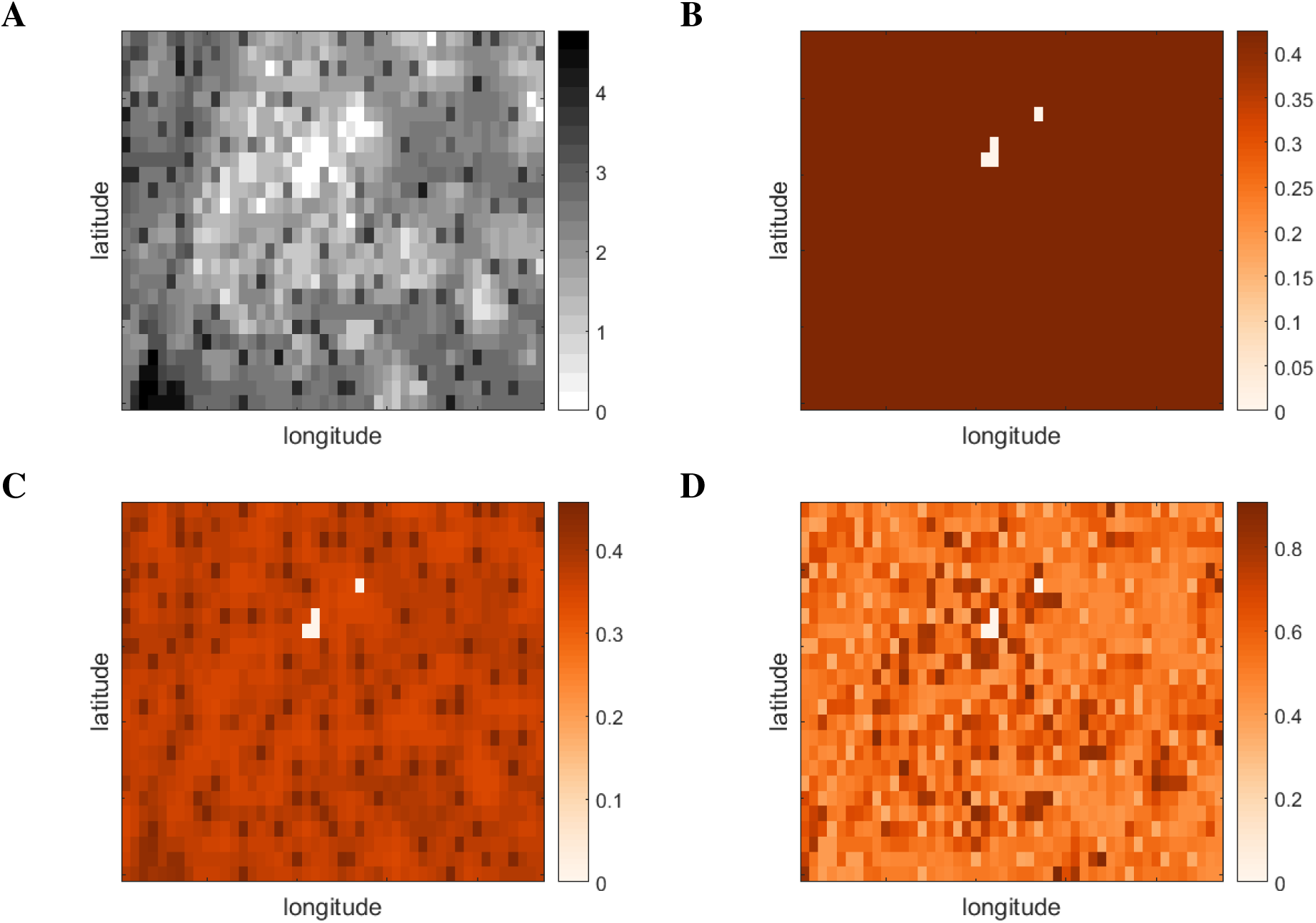
**(A)** Population density. Location-specific attack rates for **(B)** dual-mobility, **(C)** S-mobility and **(D)** I-mobility. We change colour scale between plots to better illustrate the emergent patterns. A total of 3 pixels are unpopulated and so attack rates are necessarily always zero in these locations.

Conversely, when only infectious individuals were assumed to be mobile, pixel attack rates were negatively correlated with log population density (c=−0.7707, Figure 1E) and even more strongly negatively correlated with log density gradient (c=−0.8816, Figure 1F). Attack rates varied over a greater range than for susceptible-only mobility: from a minimum of 32.61% to a maximum of 90.73%, with an absolute range of 58.12%. High attack rate pixels tended to be sparsely populated relative to neighboring locations (Figures 2A and 2D).

Measures of spatial variation are inherently dependent on the resolution of the model grid and even the strong variability outlined above would be missed by most surveillance systems. The absolute range of attack rates for the susceptible-only movement was reduced to 1.67% when aggregated to 8km by 8km pixels. Even though the effect of infectious-only movement was stronger than for susceptible-only mobility, it was rapidly hidden by the aggregation of pixels, with the absolute range dropping to 3.78% when aggregated to 8km by 8km pixels. Results of aggregation are plotted in figures S1 and S2.

The direction of association between FOI assumptions and local attack rate was preserved and the amplitude remained substantial for intermediate scenarios in which both susceptible and infectious individuals were mobile but to differing degrees. If infectious individuals had any more contacts than susceptible individuals then attack rates were negatively correlated with population density, and vice versa (Figure 3). When infectious individuals reduced their travel by a factor of 0.5, the absolute range of attack rates was 5.38% and when susceptible individuals reduced their mixing by the same degree (with infectious agents fully mobile), the absolute range was 12.89%.

**Figure 3:**
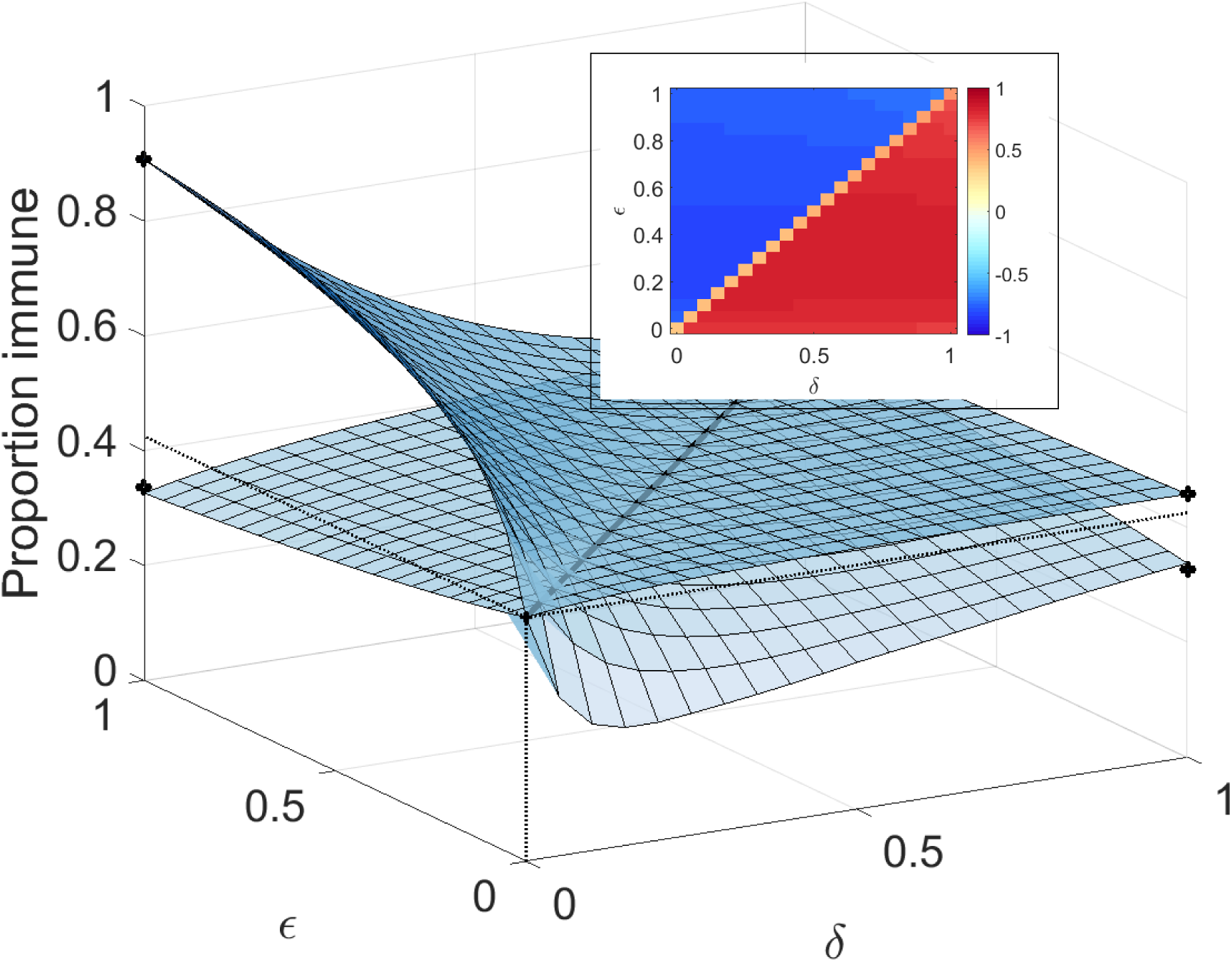
Limiting mobility of susceptible/recovered and immune agents according to parameters *δ* and *ε*, as described in the main text. Upper and lower surfaces show maximum and minimum values. Inset: correlation coefficient with population density

The underlying mobility choice kernel *K* was defined by the relative probability of making a contact in a population at a distance *x* and of population size *m*. It was parameterized by an offset distance *d*, a distance power *γ* and destination population power *α*; *K* = *m^α^/*(1 + *d/x*)^*−γ*^, with values obtained by fitting to data from this population [22]. Qualitatively, our conclusions about the impact of differential contact rates by susceptible individuals were not sensitive to values for the offset distance *d* nor the distance power *γ* (Figures 4A-4D). However, they were sensitive to values of the destination power *α* for which we have used the best fit value of 0.53 (for results up to this point) (Figures 4E, 4F). Intriguingly, with the often-assumed default value *α* = 1, the correlation between local attack rates and population density or gradient have the opposite sign (Figures S3 and S4). Moreover, *α* = 1 induces weaker correlations with local population gradient.

**Figure 4:**
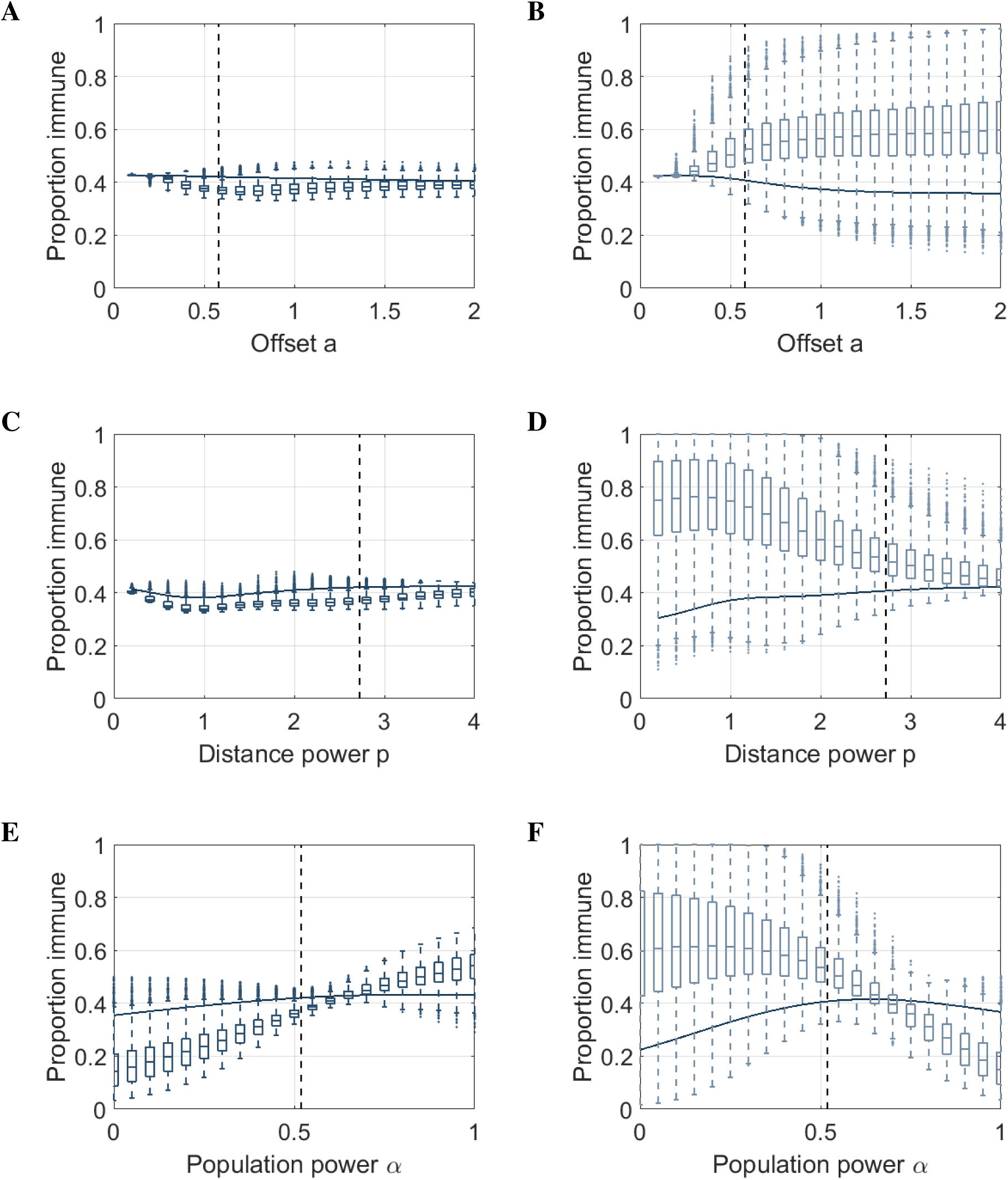
Sensitivity analysis with respect to kernel parameters *a, p* and *α* using S-mobility (LHS) and I-mobility (RHS). Box plots show standard percentiles and outliers, solid lines show global attack rate. Dual mobility are omitted as they are flat with variance *σ*^2^ = 0. Empty cells yield attack rate zero and are omitted from calculations.

Stochastic solutions to the meta-population models suggest that attack rate variation driven by asymmetric mobility would not be dominated by demographic stochasticity (Figure 5). Although attack rate variation driven may be dominated by stochastic effects for small populations, this was not the case for medium and high population densities. For pixels with the smallest population, the amplitude of variation expected to arise from asymmetric mobility is similar to that which may arise by chance due to stochastic effects. However, as the size of the pixel populations increases, the expected amplitude of stochastic variation diminishes more quickly than does the expected amplitude of variation due to asymmetric mobility (Figure S5). For example, using susceptible-only mobility for 1km by 1km pixels with populations between 1 and 85,163, the standard deviation in attack rate due to stochasticity is 9.45% while the standard deviation of expected attack rates due to asymmetric mobility is 2.61%.

**Figure 5:**
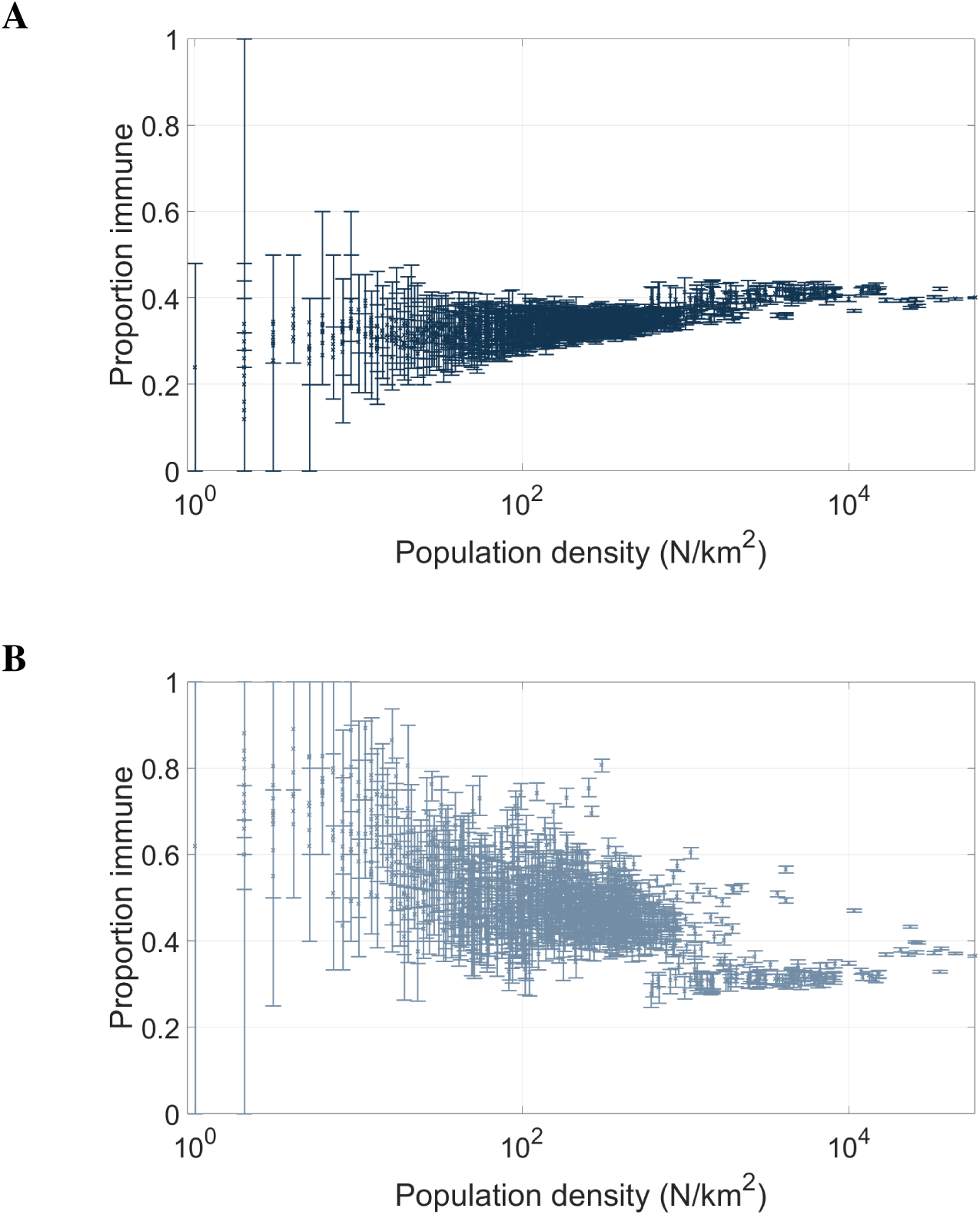
Mean and IQR over 50 iterations of stochastic model, using **(A)** S-mobility and **(B)** I-mobility.

These results are robust to our choice of illustrative population density and to alternate natural history parameters. The same effects are observed when using population density of Puerto Rico with influenza natural history parameters (Figures S6) and with parameters that approximate vector-borne transmission, such as those of Zika of Chikungunya (Figure S7). Summary statistics for these alternate scenarios for a range of deterministic model variants we have studied are shown in Table S1.

## Discussion

We have shown that, under the assumption that an individual’s total contact is independent of home location and where they travel, substantial heterogeneity in local attack rates could arise if mobility is dependent on infection status. Moreover, the direction of the relationship between attack rate and population density is dependent on the relative attractiveness of densely populated destinations compared to less dense destinations. For the best estimate of that scaling for our population of interest, when susceptible individuals are more mobile than infectious individuals, attack rates are negatively correlated with population density. Conversely, for the often implicitly assumption that the kernel is directly proportional to population size, susceptible-only movement is positively correlated with population density. Our study has a number of limitations. We have not considered spatial variation in the age distribution of people, because these data were not available for our study population. Variability in local attack rates will very likely also be driven in non-trivial ways by spatial correlation in the proportion of the population in different age classes. Nor have we considered multiple years of transmission which would extend the applicability of our results beyond pandemic scenarios for influenza and other emergent pathogens.

The negative correlation between local attack rate and population density that we observe with the most likely parameter values for some mobility assumptions is intriguing and somewhat counter intuitive. However, our sensitivity analysis with respect to kernel population power *α* provides insight into the underlying mechanisms. For example, consider the special case where only infectious people are mobile and *α* tends to large values, making mobility dependent only on population density of location, and not on geographical distance. Under this scenario, high density cells will draw in more and more infectious people and therefore generate higher attack rates. Conversely, if *α* = 0, then mobility is dependent only on distance. Under this scenario, we can think of the infectious populations spilling out of their home locations into neighbouring ones. Thus, any sparsely populated location that is adjacent to a densely populated location will see an influx of infectious individuals resulting in a greater *proportion* infectious in that location, and therefore a stronger FOI and subsequent attack rate (Figure 6).

**Figure 6:**
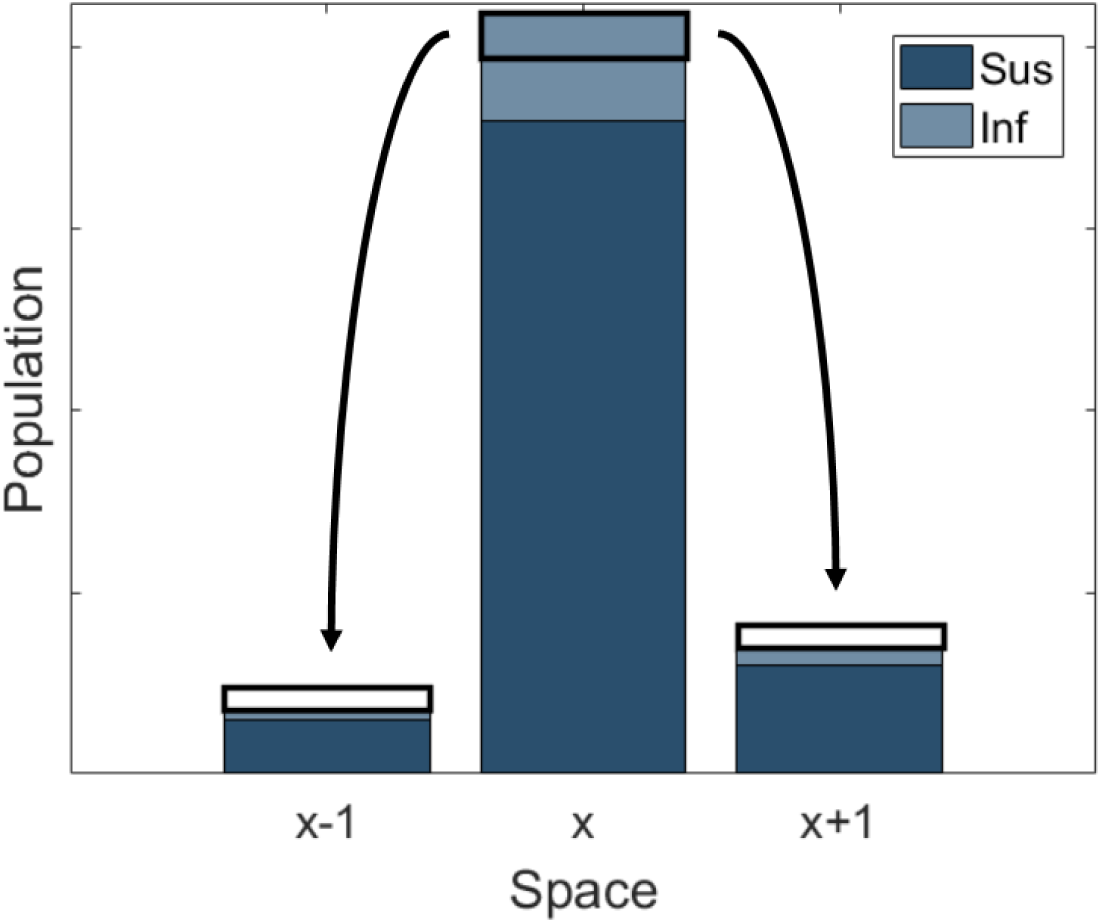
Illustration of the transmission process with infectious-only mobility: location *x* is locally densely populated, and prevalence is initially proportional to population density. If the travel kernel *K* is dominated by distance (*α* small, c.f. figure S3a), then some of the infectious population in each pixel will relocate to neighbouring pixels. The result is a higher prevalence in locally sparsely populated pixels. Moreover, a larger local gradient will allow this phenomenon to persist. Infection status is recorded by home location, which, under the I-mobility assumption, is equivalent to location when susceptible/recovered.

These results illustrate the potential knock-on effects of little or no dependence between transmissibility and population density: that infectious people from more densely populated areas go to nearby sparsely populated areas and in some sense "seek out" people in those areas to infect so they can reach their quota (I-mobility). Within the realm of parameters that are supported by studies of human movement and infectious processes, the behaviors implied by the models we presented here seem valid.

Mobility assumptions also have implications for the interpretation of attack-rates derived from individual-based models, many of which assume implicitly that the spread of infection is driven by the movement of individuals. We have shown that, whichever mobility assumption is made in a given model, it is possible to modify this assumption by replacing isotropic *K* by a convoluted non-isotropic kernel *L* that accounts for different contact patterns. In particular, the low-prevalence assumption makes this transformation achievable with minimal modification to existing computer programs. Therefore, developers of individual-based models may wish to consider alternate connectivity matrices for their simulations so as to explicitly reflect different spatial assumptions about the force of infection.

We have also shown that the implications of typical assumptions that are made in spatially explicit FOI terms, including approximations to this crucial normalization, are non-trivial at small spatial scales. Such assumptions are, however, often not addressed explicitly and so may contribute unknowingly to results. We hope to offer clarity in the interpretation of FOI in spatial models, and to have provided a comprehensive framework from which we can gain a deeper understanding of the role of spatial mobility in disease transmission dynamics as infectious disease incidence data become available at higher and higher spatial resolution.

## Methods

### Spatial kernels

Data taken from populations we study here show that total contacts made per day, and contact durations, do correlate with population density (*p <* 0.001, [11]), but that the strength of the relationship is weak. This is in part due to a typically older populations in urban areas, but also to the phenomenon of urban isolation [24]. When investigating the effect of mobility assumption alone in FOI, our main results made the baseline assumption that total contact and duration of contact is independent of home location.

The way in which these contacts are distributed in space does, however, depend on distance and population density, and is described via a spatial kernel *K*. In matrix notation, *K_ij_* is defined as the proportion of time spent by an agent from location *i* in location *j*. The assumption of uniformity of total contact therefore means that the rows of *K* sum to unity. Our model employs the offset gravity kernel, defined as follows:

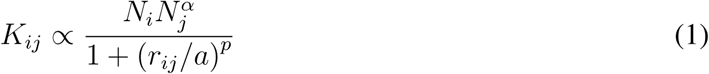

with baseline parameters of *a* = 0.58*, p* = 2.72*, α* = 0.52, where *r_ij_* denotes the geodesic distance between the centre-points of cells *i* and *j*. Of the kernel structures studied in [22], offset gravity is shown to best represent contact data. Imposing the constraint that *K* is stochastic renders redundant the factor *N_i_* in the numerator (owing to row-normalisation).

### Population density map

We used rectangular excerpts from the Landscan dataset [25] with the lower left corner of the rectangle located on the centre of the city of Guangzhou, China. The rectangle is 55 k from east to west and 33 k from north to south, and a 4 k boundary area was excluded after simulation.

The boundary area was chosen according to the following rationale: when population density data for large suburban areas is truncated for the purpose of simulation, it is equivalent to imposing empty space outside of the boundary, and this modification may effect the attack rates calculated in cells close to that boundary. We ran simulations on a large area of 1km by 1km squares, and on smaller areas contained within this larger area. We found that attack rates agree on all cells on the interior of the smaller area once a 4km perimeter is removed.

### Force-of-infection

Let *A* denote the S-mobility kernel and *B* the I-mobility kernel. Then the age-independent generalized FOI equation is given by:

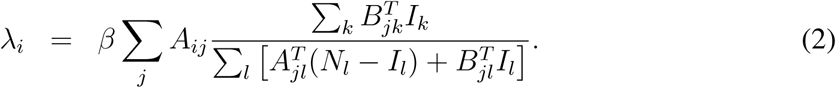

For reduced mobility, movement of the non-infectious population is governed by a parameter *δ* such that *A* = (1 *− δ*)*E* + *δK*, where *E* is an identity matrix representing absence of spatial mobility. Similarly, we describe mobility of infective individuals by *ε* such that *B* = (1 *− ε*)*E* + *εK*.

If *K* is the *n × n* spatial kernel, indexed by *i, j, k, l*, and *C* the 4 *×* 4 age-mixing matrix, indexed by *a, b, c, d*, then the age-explicit dual-mobility equation is given by:

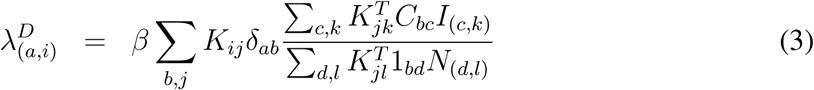

where 1_*bc*_= 1 *∀b, d*. This can be combined with equation (2) to give the age-dependent system with reduced mobility.

In all simulations presented in this study, we use the pointwise product of the matrices defining number of contacts and duration of contact between age groups 0 *−* 4, 5 *−* 19, 20 *−* 64 and 65+ derived in [11]. These age-mixing matrices were constructed from a contact surveys conducted in the region of Guangzhou used in our results.

### Model Solutions

We define the gridded transmission model as ordinary differential equations. However, we also implement a stochastic compartmental version of the model and we calculate attack rates using recursive equations.

We used a standard SIR model with 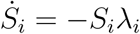,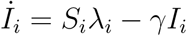,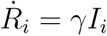. ODE models were seeded proportional to population density (*σ* = 10^*−*4^ *×* **N***/ ∑_i_N_i_*), and agreed with final size calculations (which assume infinitesimal seeding). Integration of ODEs with full FOI in the S- and I-mobility case, i.e. with *I_l_*(*t*) in denominators, showed low-prevalence approximations to be good. For example, in the main S-mobility result, the mean difference in cell attack rates between the full FOI and low prevalence approximation was 6.22 *×* 10^*−*4^ with maximum difference 3.3 *×* 10^*−*3^ occurring in a cell with population 726. Therefore, numerical solutions for all figures were obtained using the low prevalence approximation with infectious individuals in the numerator of the FOI but not the denominator (equations (18) and (22)). A selection of smaller examples agreed when checked using the full FOI.

The stochastic compartmental variant of our model selected the number of agents to infect from binomial distribution with parameters *S*_(*a,i*)_ and 1 *−* exp(*−λ*_(*a,i*)_). This method requires specification of a time-step, and we found ∆*t* = 1*/*6 days to be sufficiently small (results did not change when ∆*t* is doubled).

## Protocol S1: Algebraic analyses

### Uniform local attack rates for dual mobility assumptions

To show uniformity of attack rate with respect to space, we construct the final size equation for the system. If the final size for age-group *a* in location *i* is given by *z_a,i_* = *R_a,i_/N_i_*, then

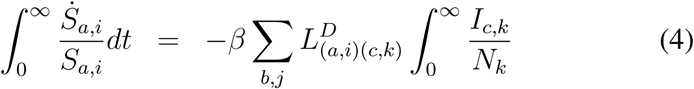

where

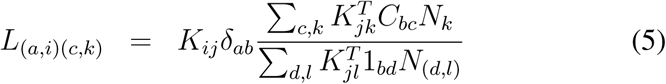

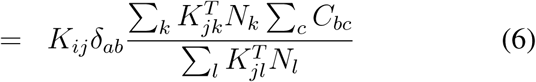

so

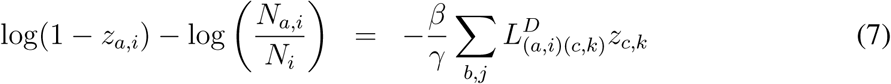

As reasoned in Ref [26] (for the S-mobility FOI with denominator *N_i_*, and in the absence of age-mixing), if there exists a solution **z** such that *z_a,i_* = *x_a_*, i.e. final sizes are independent of space, then we have:

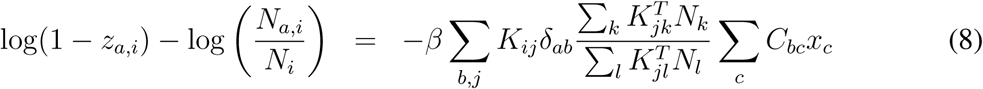

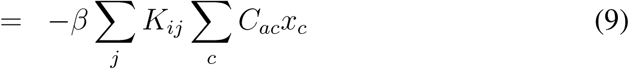

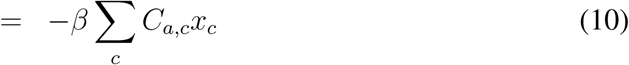

If the distribution of age-groups is uniform in space, then we have *N_a,i_/N_i_* = *q_a_*, and so, if there exists a solution to the age-only final size equation:

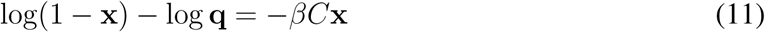

then *z_a,i_* = *x_a_* is a solution to equation (7), and final sizes are uniform in space.

### Relationship to other approximations in the literature

It is shown in [26] that susceptible-only mobility induces uniformity of attack rate, using a FOI with normalization by native population. In fact, we can show that uniformity is guaranteed only when all agents are equally mobile, owing to denominator in the force of infection term, which must be corrected to account for spatial mobility within the whole population.

For ease of notation, the following formulae are presented without explicit reference to age-mixing, but this is always included in computational results (c.f. methods for age-explicit formulae). The dual mobility FOI assumes that all agents are fully mobile as described by the kernel *K* (a stochastic matrix). The dual mobility FOI on an agent resident in pixel *i* is given by

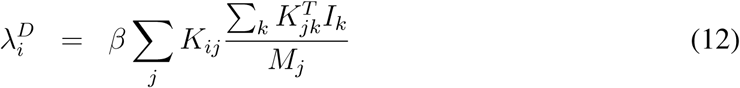

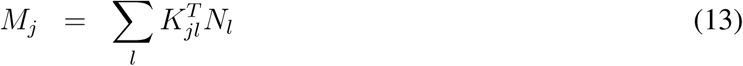

where *M_j_* denotes the total population present in cell *j*. Crucially, when a model incorporates spatial mobility, we can not say *M_j_* = *N_j_*. This FOI assumes frequency-dependent transmission based on constant contacts, and describes the expected dynamics in an agent-based system with explicit travel determined by *K*.

In the literature, the S- and I-mobility kernels are typically denoted as follows:

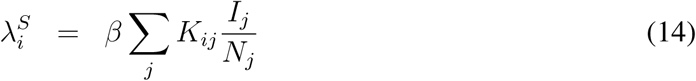

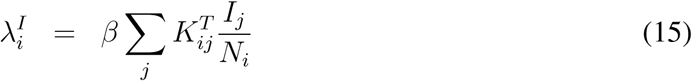

We claim that the denominators *N_j_* and *N_i_* do not accurately represent the population present in cells *j* and *i* respectively in high resolution gridded models, owing to spatial mobility. The argument below shows that the classic IM FOI serves as at least as a good approximation when incidence is small, but the SM FOI does not.

Using the above equations, SM and DM both induce uniform cumulative attack rates in space. The real SM FOI is significantly different to equation (14) in a way that is described by the ratio of total time spent in each cell and the native population of that cell.

Consider deriving the S-mobility and I-mobility FOIs from the dual mobility FOI. This involves starting with equation (12) and replacing either the single appearance of *K* or the single appearance of *K^T^* with the identity matrix (denoted *E* to avoid confusion with the *I_i_*), and adjusting denominators *M_j_* accordingly:

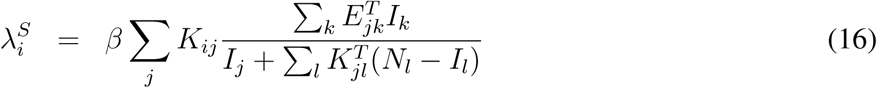

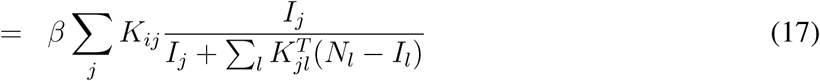

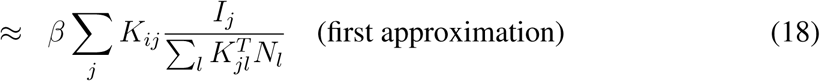

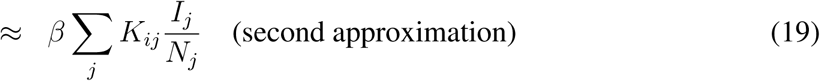

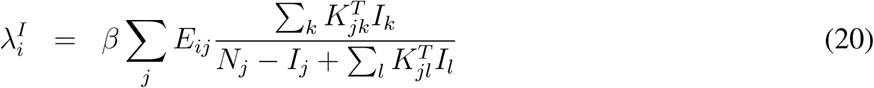

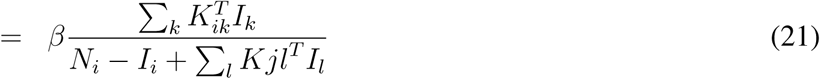

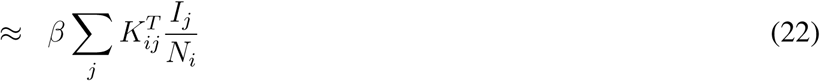

The denominator in equation (18) is not *N_j_* (the native population of cell *j*) but is instead the population *present in cell j* according to *K*. In the case where these 2 quantities are equal, we have uniformity of attack rates, as seen in the literature (using a similar argument to our proof that the DM FOI induces uniform attack rates). This approximation makes the assumption that prevalence is low, i.e. *I_i_ « N_i_ ∀i*.

The further approximation in equation (19) requires that the total number of people leaving each cell is the same as the total number of people arriving in each cell, or, because the row sums of *K* are equal to 1 (k is stochastic), then *K^T^* is also stochastic. This is a weaker assumption but is related to the *N_i_ → ∞* approximation used in [27]. When using the full FOI terms for S- and I-mobility, the only case in which these conventional mobility assumptions induce uniformity of attack rate is when each location is equally visited (in mathematical terms, uniformity of total contact means that the spatial kernel is a stochastic matrix, and the latter requirement is equivalent to the transpose of K also being stochastic, hence K is orthogonal). The notion of normalization by total population present is not new to the literature [28], though is often excluded in the construction of spatial epidemic models.

### Non-isotropic convoluted kernels

It is possible to change the mobility assumption in an existing model via an effective, or convoluted kernel *L* such that replacing *K* with *L* in a given explicit FOI is equivalent to a change of mobility assumption. In fact, we can write any spatially explicit FOI in the form:

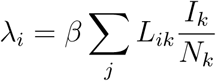

for some matrix *L*. This formulation is essential in final size calculations. Then, for example, the convoluted D-mobility kernel *L^D^* is given by 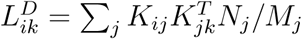, where *M_j_* = ∑_*l*_*K*_*jl*_*N*_*l*_, as in the main text. When using low-prevalence approximations, this can be done prior to numerical simulation, and so requires minimal additional modification to existing model codes.

### Global transmissibility coefficient

In all simulations, we use the next generation matrix (NGM) method [23] to derive a global transmissibility parameter *β* that yields our desired global *R*_0_. NGMs are derived from *λ_i_*, evaluated at disease-free equilibrium (DFE). We can show that, in all 3 cases, using the approximations to S- and I-mobility given in equations (18) and (22) the value of *β* obtained is equal to that of the spatially heterogeneous system, maintaining heterogeneity in age only.

In the I-mobility case, the NGM is given by

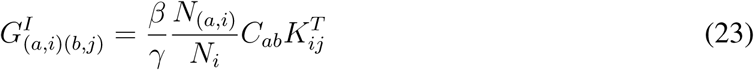

Since *K* is a stochastic matrix, we have *λ*_1_(*K*) = 1 and so *λ*_1_(*K^T^*) = 1, thus the dominant eigenvalue of *G^I^* is equal to the dominant eigenvalue of

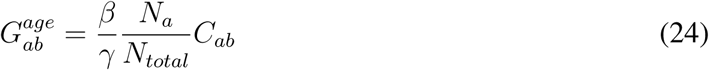

Using S-mobility, we have

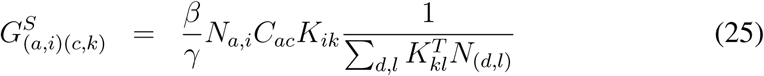

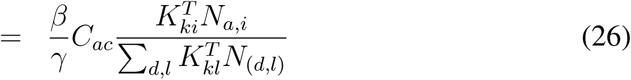

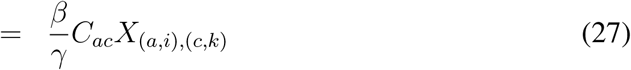

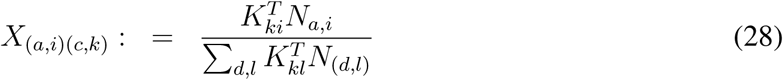

Note that *X* is a stochastic matrix, and so the dominant eigenvalue of *G^S^* is equal to the dominant eigenvalue of *G^age^*.

A similar argument applies to dual mobility, where we have

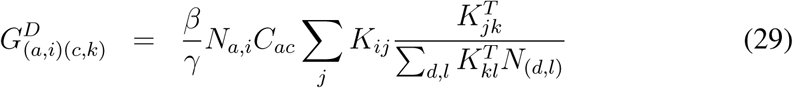

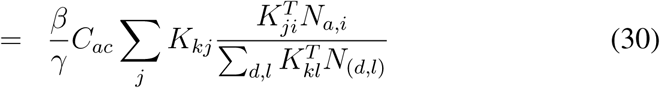

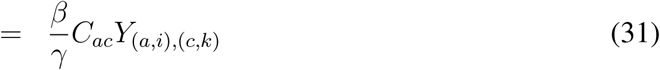

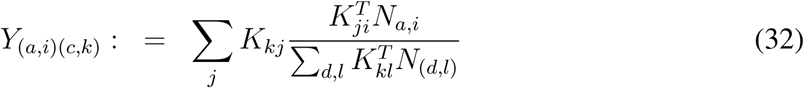

Here, since *Y* is the product of 2 stochastic matrices, it must itself be stochastic, and so the dominant eigenvalue of *G^D^* is also equal to the dominant eigenvalue of *G^age^*.

The arguments presented above for susceptible-only and dual mobility require that the same travel kernel *K* be used to describe the movement of all age groups, i.e. *K_ij_* be independent of *a, b, c, d*. It can be verified computationally that age-dependent mobility can indeed induce different values of *β* to the spatially heterogeneous model, in all cases other than pure infectious-only mobility. We reserve a detailed analysis of this scenario for a subsequent study.

---

**Figure S1:**
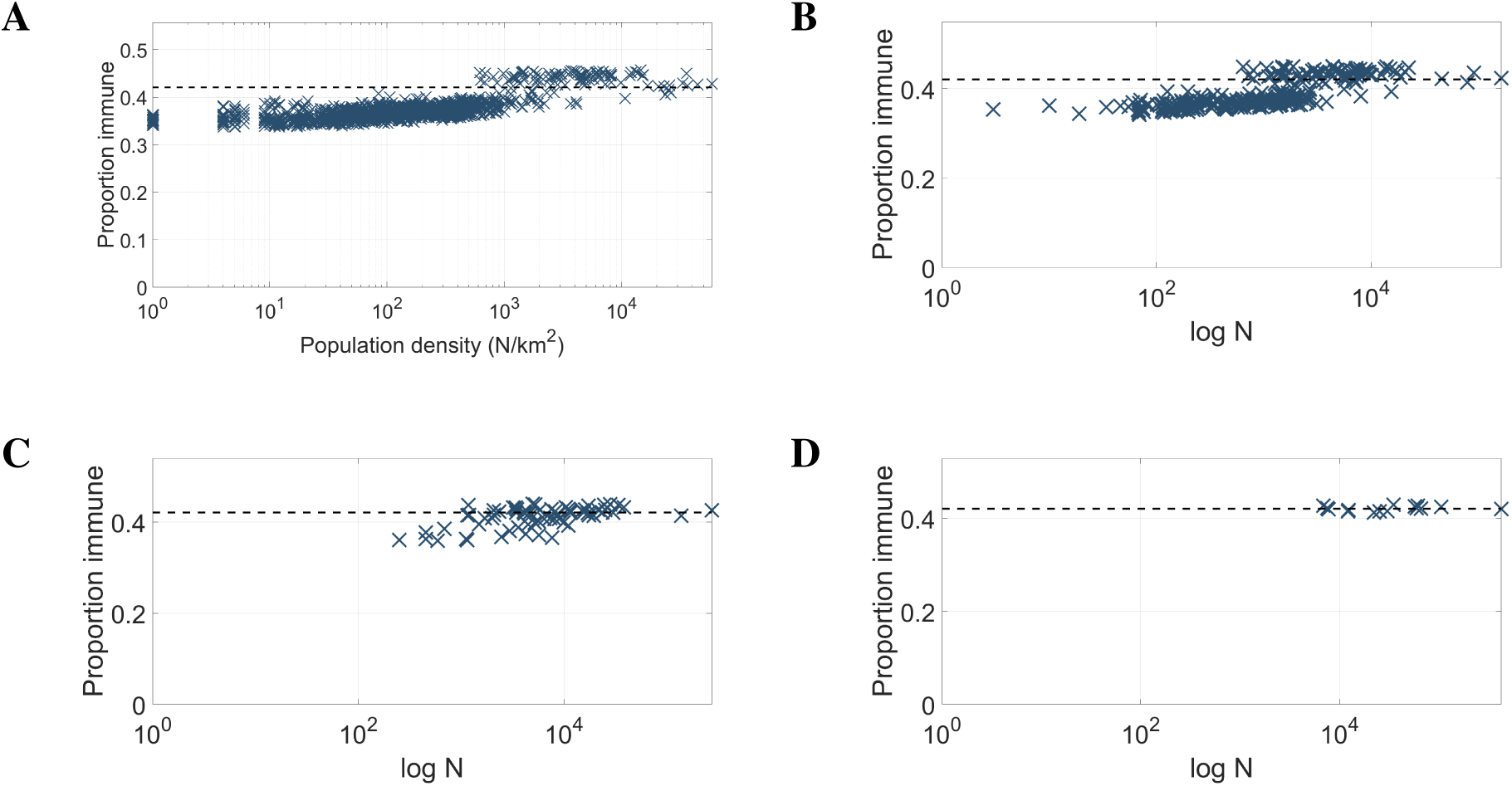
S-mobility: **(A)** initial result, aggregated into **(B)** 2km by 2km, **(C)** 4km by 4km, and **(D)** 8km by 8km squares.

**Figure S2:**
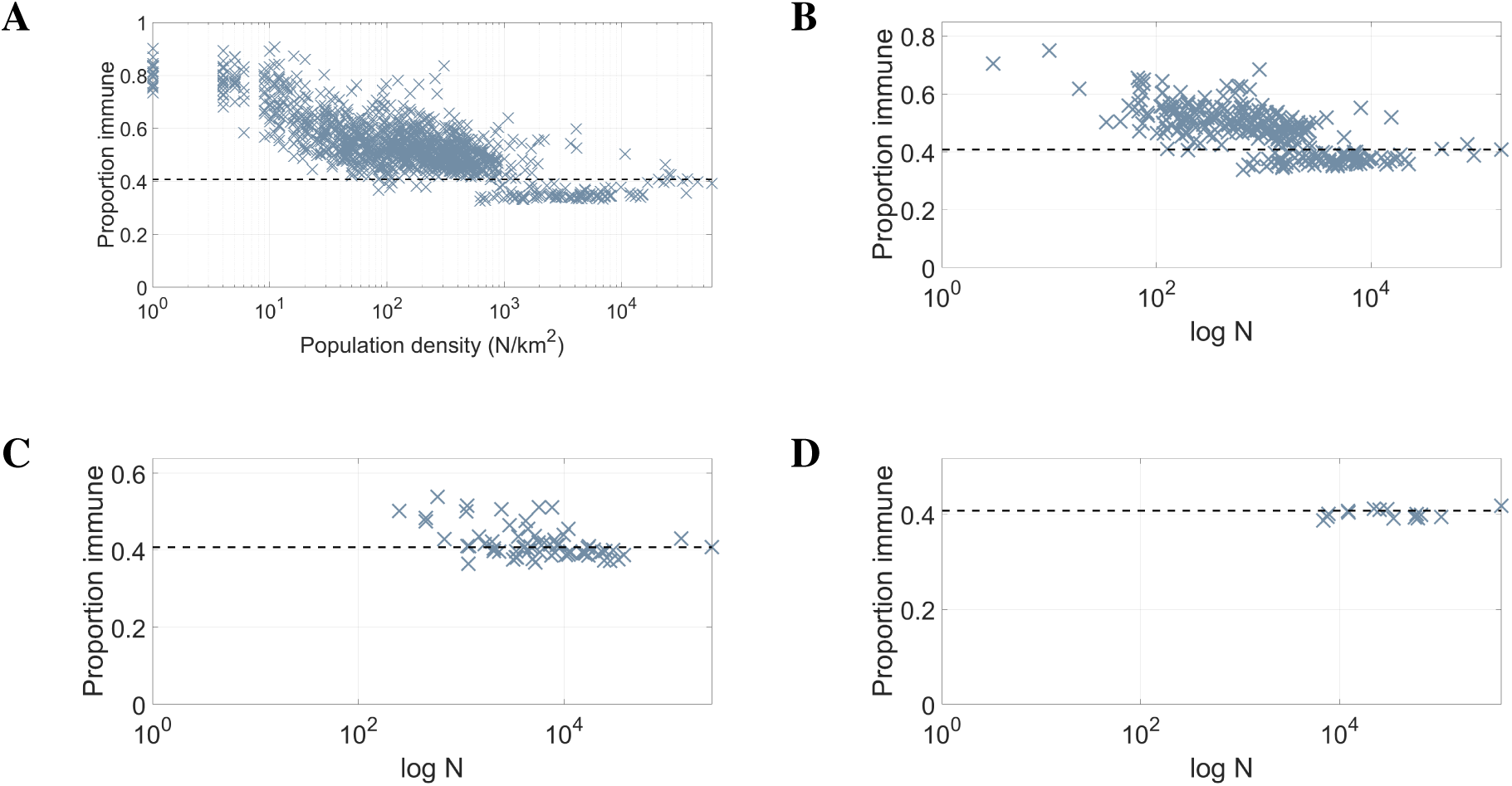
I-mobility: **(A)** initial result, aggregated into **(B)** 2km by 2km, **(C)** 4km by 4km, and **(D)** 8km by 8km squares.

**Figure S3:**
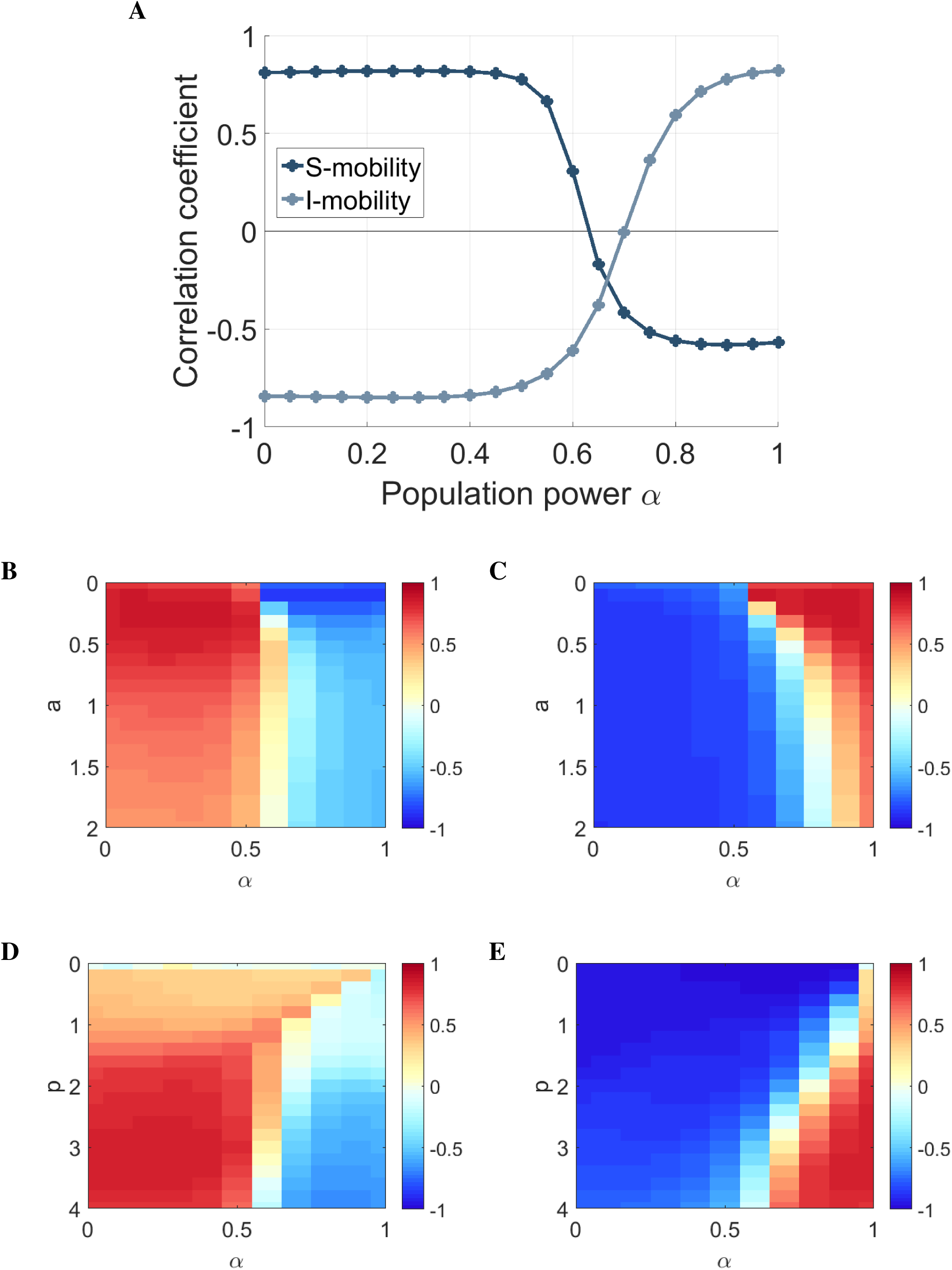
Sensitivity analysis: correlation coefficient of attack rate with population density **(A)** with *a* and *p* fixed, **(B)** S-mobility, *p* fixed, **(C)** I-mobility, *p* fixed, **(D)** S-mobility, *a* fixed, and **(E)** I-mobility, *a* fixed. Fixed parameters are set at optimal values discussed in the main text.

**Figure S4:**
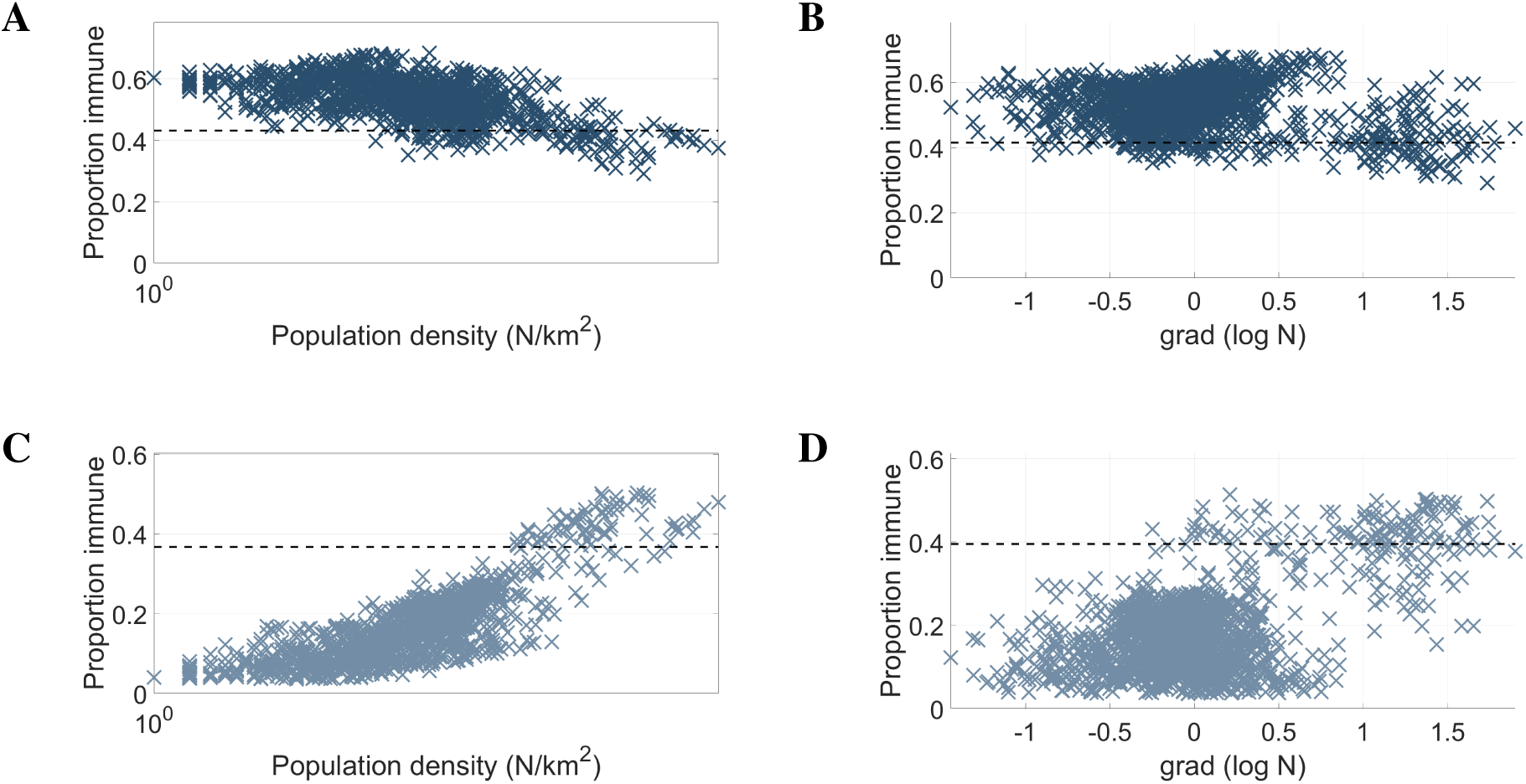
Repeating results of figure 1 with *α* = 1: **(A)** S-mobility/density, **(B)** S-mobility/gradient, **(C)** I-mobility/density, and **(D)** I-mobility/gradient.

**Figure S5:**
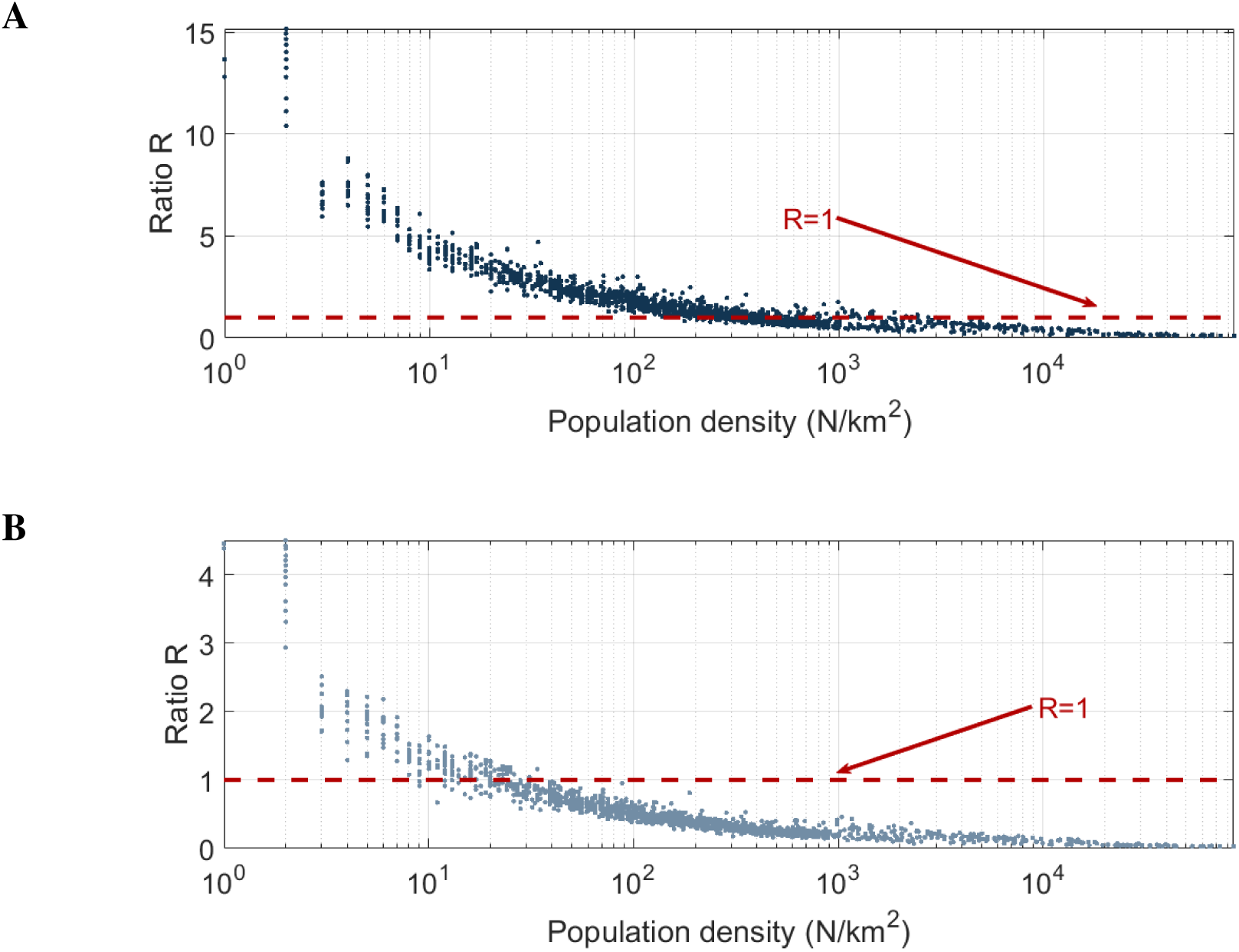
Ratio *R* of location-specific standard deviation over 50 iterations of stochastic model to standard deviation of corresponding deterministic model result over all pixels, using (a) S-mobility and (b) I-mobility.

**Figure S6:**
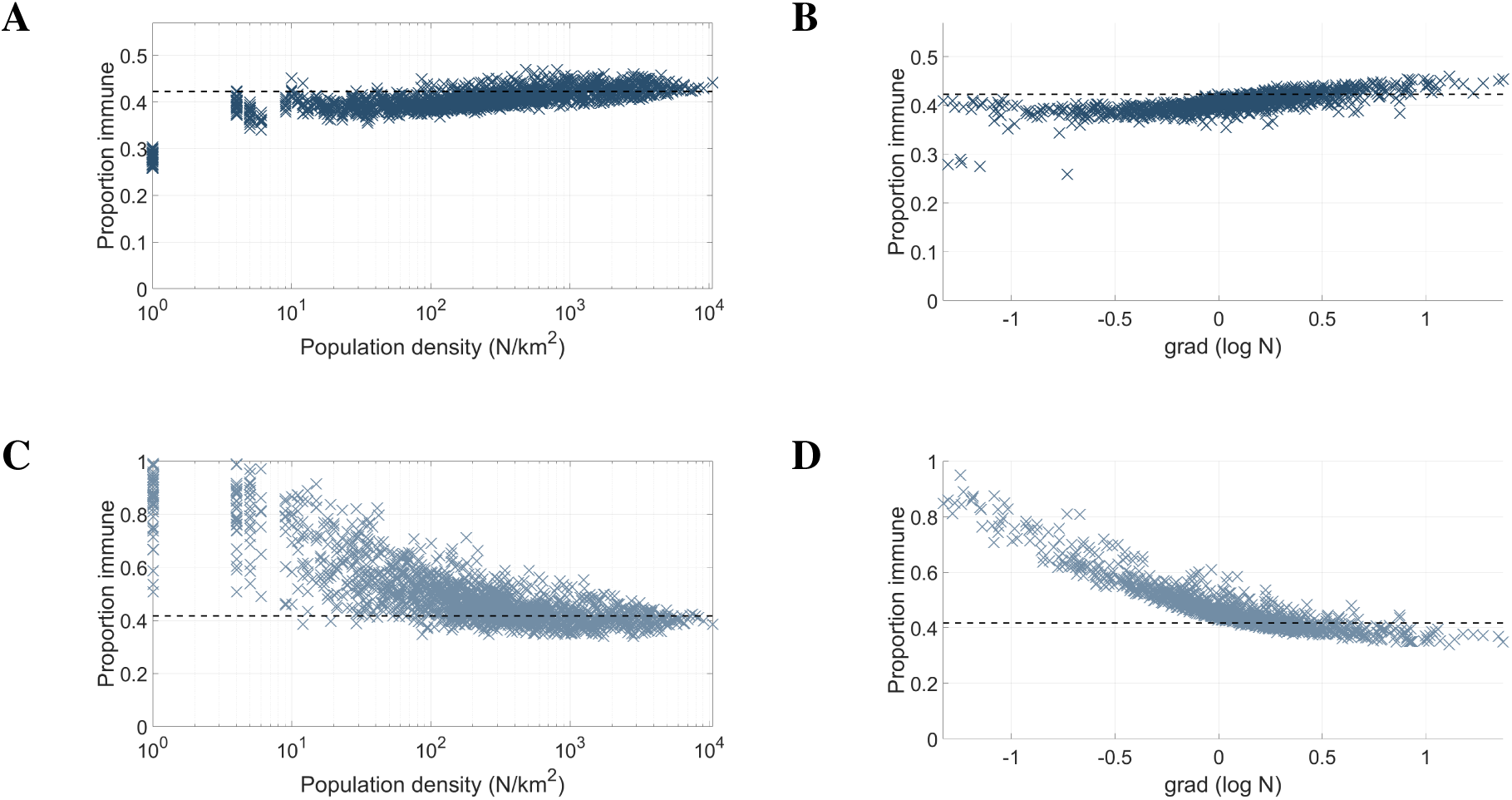
Simulated attack rates using population density of North-East Puerto-Rico, a 60km by 60km grid, and influenza-like natural history parameters *R*_0_ = 1.8*, γ* = 1*/*2.6, with **(A)** S-mobility plotted against population density, **(B)** S-mobility plotted against log population gradient, **(C)** I-mobility/density, and **(D)** I-mobility/gradient.

**Figure S7:**
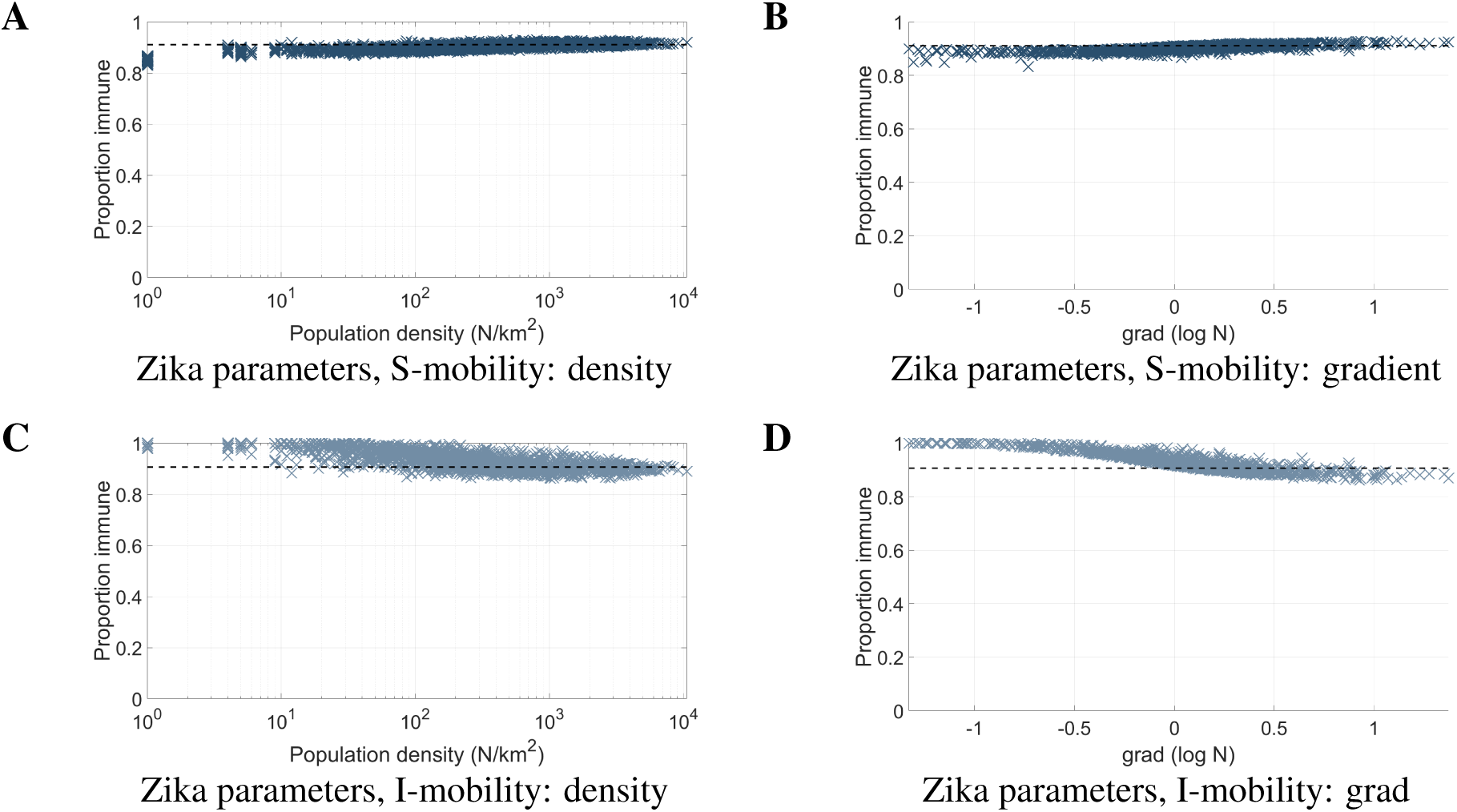
Simulated attack rates using population density of North-East Puerto-Rico, a 60km by 60km grid, and natural history parameters *R*_0_ = 4*, γ* = 1*/*10 approximating vector-borne transmission (e.g. Zika, Chikungunya), with **(A)** S-mobility plotted against population density, **(B)** S-mobility plotted against log population gradient, **(C)** I-mobility/density, and (d) I-mobility/gradient.

**Table S1:** Summary statistics for different model parameters, populations and mobility assumptions. Results for different grid sizes involve aggregation of result obtained at 1km by 1km resolution. In all cases, empty cells are omitted from calculations. It is therefore possible to obtain a smaller minimum value of attack rate after aggregation.

| Model |  |  |  | Attack rate |  |  |  |  |
| --- | --- | --- | --- | --- | --- | --- | --- | --- |
| $(R_0, \gamma, \alpha)$ | Population | Grid | Mobility ass. | Min. | Max. | Global | $CC_{pop}$ | $CC_{grad}$ |
| (1.8, 1/2.6, 0.52) | Guangzhou | 1km by 1km | DM | 0.425 | 0.425 | 0.425 | 0 | 0 |
|  |  |  | SM | 0.3372 | 0.421 | 0.4576 | 0.7469 | 0.7347 |
|  |  |  | IM | 0.3261 | 0.9073 | 0.4077 | 0.7707 | 0.8816 |
|  |  | 4km by 4km | DM | 0.4576 | 0.4576 | 0.4576 | 0 | 0 |
|  |  |  | SM | 0.5523 | 0.7259 | 0.359 | 0.4214 | 0.4399 |
|  |  |  | IM | 0.3648 | 0.5384 | 0.4076 | -0.4651 | -0.6883 |
| (1.8, 1/2.6, 1) | Guangzhou | 1km by 1km | DM | 0.425 | 0.425 | 0.425 | 0 | 0 |
|  |  |  | SM | 0.2909 | 0.6842 | 0.4316 | -0.5687 | -0.149 |
|  |  |  | IM | 0.0355 | 0.5029 | 0.3672 | 0.8213 | 0.5454 |
|  |  | 4km by 4km | DM | 0.4576 | 0.4576 | 0.4576 | 0 | 0 |
|  |  |  | SM | 0.318 | 0.6657 | 0.431 | 0.7668 | 0.768 |
|  |  |  | IM | -0.0415 | 0.5029 | 0.3687 | 0.5384 | 0.4076 |
| (1.8, 1/2.6, .052) | Puerto Rico | 1km by 1km | DM | 0.4251 | 0.4251 | 0.4251 | 0 | 0 |
|  |  |  | SM | 0.3598 | 0.4703 | 0.4231 | 0.6323 | 0.738 |
|  |  |  | IM | 0.3405 | 0.9935 | 0.4181 | -0.7915 | -0.8887 |
|  |  | 4km by 4km | DM | 0.4251 | 0.4251 | 0.4251 | 0 | 0 |
|  |  |  | SM | 0.3869 | 0.4591 | 0.4231 | 0.8866 | 0.5259 |
|  |  |  | IM | 0.3709 | 0.5932 | 0.4181 | 0.8193 | -0.8887 |
| (4, 1/10, 0.52) | Puerto Rico | 1km by 1km | DM | 0.912 | 0.912 | 0.912 | 0 | 0 |
|  |  |  | SM | 0.8683 | 0.9298 | 0.9111 | 0.6923 | 0.7631 |
|  |  |  | IM | 0.8615 | 1 | 0.9072 | -0.7528 | -0.9008 |
|  |  | 4km by 4km | DM | 0.912 | 0.912 | 0.912 | 0 | 0 |
|  |  |  | SM | 0.8851 | 0.9248 | 0.9111 | 0.8805 | 0.4554 |
|  |  |  | IM | 0.8809 | 0.98 | 0.9072 | 0.864 | -0.2702 |

## References

[1] Hay SI, Snow RW. The malaria Atlas Project: developing global maps of malaria risk. PLoS Med. 2006 Dec;3(12):e473.

[2] Bhatt S, Weiss DJ, Cameron E, Bisanzio D, Mappin B, Dalrymple U, et al. The effect of malaria control on Plasmodium falciparum in Africa between 2000 and 2015. Nature. 2015 Oct;526(7572):207–211.

[3] Charu V, Zeger S, Gog J, Bjørnstad ON, Kissler S, Simonsen L, et al. Human mobility and the spatial transmission of influenza in the United States. PLoS Comput Biol. 2017 Feb;13(2):e1005382.

[4] Lessler J, Azman AS, McKay HS, Moore SM. What is a Hotspot Anyway? Am J Trop Med Hyg. 2017 Jun;96(6):1270–1273.

[5] Sutton P, Elvidge C, Obremski T. Building and Evaluating Models to Estimate Ambient Population Denstiy. Photogrammetric Engineering and Remote Sensing. 2003;69(5):545–553.

[6] Deville P, Linard C, Martin S, Gilbert M, Stevens FR, Gaughan AE, et al. Dynamic population mapping using mobile phone data. Proceedings of the National Academy of Sciences. 2014;111(45):15888–15893. Available from: http://www.pnas.org/lookup/doi/10.1073/pnas.1408439111.

[7] Hay SI, Snow RW. The Malaria Atlas Project: Developing global maps of malaria risk. PLoS Medicine. 2006;3(12):2204–2208.

[8] Stanaway JD, Flaxman AD, Naghavi M, Fitzmaurice C, Vos T, Abubakar I, et al. The global burden of viral hepatitis from 1990 to 2013: findings from the Global Burden of Disease Study 2013. The Lancet. 2016;388(10049):1081–1088.

[9] Rey J, Stanaway D, Shepard DS, Undurraga EA, Halasa YA, Coff LE, et al. The global burden of dengue: an analysis from the Global Burden of Disease Study 2013. WwwThelancetCom/Infection. 2016;16(6):712–723.

[10] Keeling MJ, Danon L, Ford AP, House T, Jewell CP, Roberts GO, et al. Networks and the epidemiology of infectious disease. Interdisciplinary Perspectives on Infectious Diseases. 2011;2011.

[11] Read JM, Lessler J, Riley S, Wang S, Tan LJ, Kwok KO, et al. Social mixing patterns in rural and urban areas of southern China. Proceedings of the Royal Society of London B: Biological Sciences. 2014;281(1785).

[12] Gonzalez MC, Hidalgo CA, Barabasi AL. Understanding individual human mobility patterns. 2008;453(June). Available from: http://arxiv.org/abs/0806.1256{\%}0Ahttp://dx.doi.org/10.1038/nature06958.

[13] Vazquez-Prokopec GM, Bisanzio D, Stoddard ST, Paz-Soldan V, Morrison AC, Elder JP, et al. Using GPS Technology to Quantify Human Mobility, Dynamic Contacts and Infectious Disease Dynamics in a Resource-Poor Urban Environment. PLoS ONE. 2013;8(4):1–10.

[14] Perkins TA, Garcia AJ, Paz-Soldan VA, Stoddard ST, Reiner RC, Vazquez-Prokopec G, et al. Theory and data for simulating fine-scale human movement in an urban environment. Journal of The Royal Society Interface. 2014;11(99):20140642–20140642. Available from: http://rsif.royalsocietypublishing.org/cgi/doi/10.1098/rsif.2014.0642.

[15] Quick J, Grubaugh ND, Pullan ST, Claro IM, Smith AD, Gangavarapu K, et al. Multiplex PCR method for MinION and Illumina sequencing of Zika and other virus genomes directly from clinical samples. Nat Protoc. 2017 Jun;12(6):1261–1276.

[16] Paul P, Heng BH, Seow E, Molina J, Tay SY. Predictors of frequent attenders of emergency department at an acute general hospital in Singapore. Emerg Med J. 2010 Nov;27(11):843–848.

[17] Funk S, Bansal S, Bauch CT, Eames KTD, Edmunds WJ, Galvani AP, et al. Nine challenges in incorporating the dynamics of behaviour in infectious diseases models. Epidemics. 2015;10:21–25. Available from: http://dx.doi.org/10.1016/j.epidem.2014.09.005.

[18] Funk S, Salathé M, Jansen VAA. Modelling the influence of human behaviour on the spread of infectious diseases: a review. J R Soc Interface. 2010 Sep;7(50):1247–1256.

[19] Campbell GL, Hughes JM. Plague in India: a new warning from an old nemesis. Ann Intern Med. 1995 Jan;122(2):151–153.

[20] Riley S. Models of Infectious Disease. Science. 2007;316(5829):1298–1301. Available from: http://www.sciencemag.org/cgi/content/abstract/316/5829/1298.

[21] Riley S, Eames K, Isham V, Mollison D, Trapman P. Five challenges for spatial epidemic models. Epidemics. 2015;10:68–71. Available from: http://dx.doi.org/10.1016/j.epidem.2014.07.001.

[22] Read J, Mills H. Is this on biorXiv yet?; 2017.

[23] Diekmann O, Heesterbeek JAP. Mathematical Epidemiology of Infectious Diseases: Model Building, Analysis and Interpretation. 1st ed. Wiley; 2000.

[24] Kinenberg E. Dying Alone; 2001.

[25] Dobson JE, Bright EA, Coleman PR, Durfee RC, Worley BA. LandScan: A global population database for estimating populations at risk. Photogrammetric Engineering and Remote Sensing. 2000;66(7):849–857. Available from: http://apps.webofknowledge.com.libezp.lib.lsu.edu/full_record.do?product=UA&search_mode=GeneralSearch&qid=3&SID=1D8vegIjNxsjNKm4nMs&page=1&doc=3.

[26] Ma J, Earn DJD. Generality of the final size formula for an epidemic of a newly invading infectious disease. Bulletin of Mathematical Biology. 2006;68(3):679–702.

[27] Clancy D, Pearce CJ. The effect of population heterogeneities upon spread of infection. Journal of Mathematical Biology. 2013;67(4):963–987.

[28] Sattenspiel L, Dietz K, Sattenspiel, L KD. A structured epidemic model incorporating geographic-mobility among regions. Mathematical Biosciences. 1995;128(1-2):71–91. Available from: http://ac.els-cdn.com/002555649400068B/1-s2.0-002555649400068B-main.pdf?{\_}tid=ff279706-2cec-11e4-969f-00000aab0f27{\&}acdnat=1409035893{\_}54eb62e0eb7fc6ef3e78e7374f1d5343.

